# Deconvolution of plasma pharmacokinetics from dynamic heart imaging data obtained by SPECT/CT imaging

**DOI:** 10.1101/2022.11.17.517003

**Authors:** Zengtao Wang, Lushan Wang, Malik Ebbini, Geoffry L. Curran, Paul H. Min, Ronald A. Siegel, Val J. Lowe, Karunya K. Kandimalla

## Abstract

Plasma pharmacokinetic (PK) data is required as an input function for graphical analysis (e.g., Patlak plot) of single positron emission computed tomography/computed tomography (SPECT/CT) and positron emission tomography/CT (PET/CT) data to evaluate tissue influx rate of radiotracers. Dynamic heart imaging data is often used as a surrogate of plasma PK. However, accumulation of radiolabel (representing both intact and degraded tracer) in the heart tissue may interfere with accurate prediction of plasma PK from the heart data. Therefore, we developed a compartmental model, which involves forcing functions to describe intact and degraded radiolabeled proteins in plasma and their accumulation in heart tissue, to deconvolve plasma PK of ^125^I-amyloid beta 40 (^125^I-Aβ_40_) and ^125^I-insulin from their dynamic heart imaging data. The three-compartment model was shown to adequately describe the plasma concentration-time profile of intact/degraded proteins and the heart radioactivity time data obtained from SPECT/CT imaging for both tracers. The model was successfully applied to deconvolve the plasma PK of both tracers from their naïve datasets of dynamic heart imaging. In agreement with our previous observations made by conventional serial plasma sampling, the deconvolved plasma PK of ^125^I-Aβ_40_ and ^125^I-insulin in young mice exhibited lower area under the curve (AUC) than the aged mice. Further, Patlak plot parameters (Ki) extracted using deconvolved plasma PK as input function successfully recapitulated age-dependent blood-to-brain influx kinetics changes for both ^125^I-Aβ_40_ and ^125^I-insulin. Therefore, the compartment model developed in this study provides a novel approach to deconvolve plasma PK of radiotracers from their noninvasive dynamic heart imaging. This method facilitates the application of preclinical SPECT or PET imaging data to characterize distribution kinetics of tracers where simultaneous plasma sampling is not feasible.

## Introduction

In nuclear medicine, plasma PK is required as an input function in graphic analyses such as Patlak plots to evaluate the influx kinetics of radiotracers from plasma to peripheral tissue (Patlak, Blasberg et al. 1983, Patlak and Blasberg 1985). In our previous publications, we used Patlak plots to calculate brain influx rates of ^125^I-insulin and ^125^I-amyloid beta (^125^I-Aβ) by analyzing single positron emission computed tomography/computed tomography (SPECT/CT) dynamic imaging data (Swaminathan, Ahlschwede et al. 2018, Sharda, Ahlschwede et al. 2021). However, in these studies, plasma PK and imaging studies were conducted on separate cohorts of animals due to the infeasibility of plasma sampling during in vivo imaging. Limitations of this method are 1) additional reagent, animal, and labor costs; and 2) biased estimation of the Patlak plot slope, which denotes the blood-to-brain influx clearance (Ki). It is imperative to develop novel, low-cost methods to obtain both plasma PK and tissue distribution kinetics information from the same subject.

Dynamic heart imaging data has been reported as a surrogate for plasma PK and used as the input function for kinetic analysis of various tracers including small-molecule ^18^F-FHBG (Green, Nguyen et al. 2004), and large molecule fluorescent antibody-drug conjugates (Giddabasappa, Gupta et al. 2016). Recent studies conducted by Bao et.al compared noninvasive heart fluorescence molecular tomography (FMT) and serial blood sampling to determine blood PK of probes ranging from 2-150 kDa. They found excellent agreement in kinetic profiles and associated PK parameters between these two methods and further applied FMT to estimate glomerular filtration rate under various pathological conditions (Bao, Vasquez et al. 2019). However, for molecules such as insulin and Aβ, which have been the focus of research in our lab, this method can overpredict plasma levels because both proteins, especially Aβ, exhibit considerable accumulation in heart tissue (Troncone, Luciani et al. 2016, Martinez-Naharro, Hawkins et al. 2018). The imaging signal from heart includes not only intravascular tracers but also those in extravascular space. Accordingly, any later analysis could be erroneous if the longitudinal heart dataset is used as a replacement for plasma PK.

In order to resolve this issue, we have separated the signal coming from blood and extravascular space by constructing a three component PK model and successfully deconvolved the plasma PK from dynamic heart imaging data. The model was calibrated using plasma concentration-time and dynamic heart imaging data for two different tracers, ^125^I-Aβ_40_ and ^125^I-insulin. Further, the calibrated model was applied to naïve datasets to derive plasma PK parameters and generate Patlak plots determining brain influx rate of these two tracers. The advantage of this kinetic modeling approach is the ability to infer plasma PK parameters from dynamic heart imaging data, thus eliminating the need for additional plasma PK studies. Moreover, heart dynamic imaging data is obtained from the same subjects in which brain imaging is performed, thereby yielding more accurate estimation of blood-to-brain influx rates.

## Materials and Methods

### Materials

Aβ_40_ peptide was procured from AAPPTec, LLC (Louisville, KY). Carrier-free Na^125^I radionuclides were obtained from Perkin Elmer Life and Analytical Sciences (Boston, MA). Insulin (Novolin^®^ or Humulin^®^) was purchased from Eli Lilly (Indianapolis, IN).

### Animals

B6SJLF1/J mice were purchased from Jackson Laboratory (Barbor, ME). Mice were housed in the Mayo Clinic animal care facility with food and water ad libitum under 12 h light and dark cycles. Experiments were conducted in adherence with the guide for care and use of laboratory animals provided by the National Institutes of Health. All protocols were approved by the Mayo Clinic Institutional Animal Care and Use Committee.

### Radioiodination of insulin and Aβ_40_

Insulin and Aβ_40_ were radiolabeled with ^125^I by the chloramine-T procedure as described in our previous publications (Kandimalla, Curran et al. 2005). Free ^125^I was removed from the radiolabeled protein by dialysis overnight in 0.01M phosphate-buffered saline (PBS). Purity of the ^125^I-labeled proteins was determined by trichloroacetic acid (TCA) precipitation. Specific activities of ^125^I-insulin and ^125^I-Aβ_40_ were determined to be in the range 45−48 μCi/μg.

### Plasma pharmacokinetics of ^125^I-Insulin and ^125^I-Aβ_40_

The femoral vein and artery of each animal were catheterized under general anesthesia with a mixture of isoflurane (1.5%) and oxygen (4 L/min). A single intravenous injection (100 μCi) of ^125^I-insulin or ^125^I-Aβ_40_ was administered in the femoral vein, and blood (20 μL) was sampled from the femoral artery using heparinized capillary tubes at various time points. Blood samples were diluted with saline and centrifuged to separate the plasma. TCA precipitation of plasma was conducted to separate intact and degraded ^125^I-insulin or ^125^I-Aβ_40._ The supernatant consisted of degraded protein while the precipitate was assumed to contain intact protein. Total radioactivity counts in both supernatant and precipitate were analyzed using a gamma counter (Cobra II; Amersham Biosciences Inc., Piscataway, NJ). A two-compartment model was fitted to plasma concentration-time data of intact proteins using Phoenix WinNonLin_®_ Version 8.3 (Certara USA, Inc., Princeton, NJ). Plasma concentration-time data of both degraded tracers were fitted by empirical functions using GraphPad Prism 9.1 (GraphPad software; La Jolla, CA).

### Dynamic SPECT/CT studies of _125_I-Insulin and _125_I-Aβ_40_

To assess dynamic radioactivity changes in heart and brain, mice were administered 500 μCi of ^125^I-insulin (n=8) or ^125^I-Aβ_40_ (n=9) through the femoral vein. The animals were immediately imaged for the following 42 min at 1-min intervals by dynamic SPECT/CT (Gamma Medica, Northridge, CA) as described previously (Jaruszewski, Curran et al. 2014). Both CT and SPECT images were reconstructed once the scan was acquired. A Feldkamp reconstruction algorithm was used for CT reconstruction whereas the 3-D reprojection reconstruction (3DRP) method was used for SPECT images. The reconstructed images were then loaded into PMOD software (PMOD Technologies, Switzerland), regions of interest (ROIs) was drawn using CT images.

### Determination of heart cavity volume

The volume of the heart cavity (V_h_) was calculated separately for ^125^I-insulin and ^125^I-Aβ_40_ by the following equation,

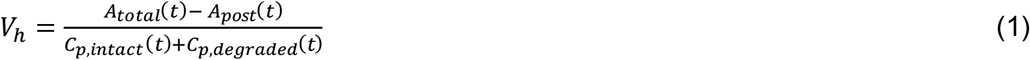

where *A*_*total*_(*t*) is the total heart radioactivity at the end of imaging (t = 42 min); A_post_(t) is the total heart radioactivity after transcardial perfusion; *C*_*p,intact*_(*t*) and *C*_*p,degraded*_(*t*) are the predicted plasma concentration of intact and degraded proteins at 42 min, respectively.

### Model development

A three-compartment model, depicted in **Figure 2**, was constructed including intact radiolabeled protein (compartment 1), degraded radiolabeled protein (compartment 2) in plasma, and the accumulation of radiolabel in heart tissue (compartment 3). Plasma concentrations of intact proteins followed biexponential decay kinetics, and the heart radioactivity, *A*_1_ (*t*), was described by

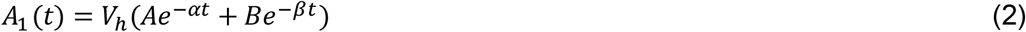

Plasma radioactivity associated with degraded protein in the heart cavity, *A*_2_(*t*), was described by

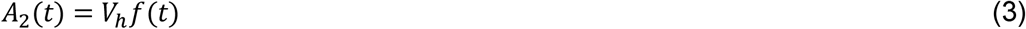

where *f*(*t*) is an empirical function that depends on the protein of interest:

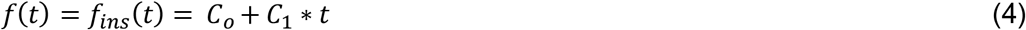

for degraded ^125^I-insulin and

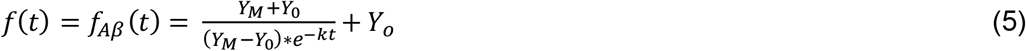

for degraded ^125^I-Aβ_40_.

The amount of radioactivity in heart tissue, *A*_3_(*t*), was described using a one-compartment model with the two input rates, *k*_1_*A*_1_(*t*) and *k*_2_*A*_2_(*t*), representing transfer of radioactivity from plasma, and *k*_0_*A*_3_(*t*), representing first order transfer out of the tissue. Thus,

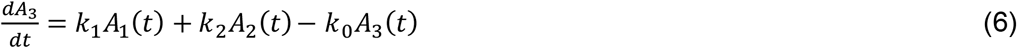

Total heart radioactivity, *A*_*tot*_(*t*), was represented by the summation of three components.

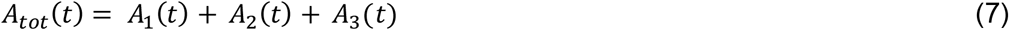

The parameters *A, α*, B, *β, C*_*0*_, *C*_*1*_, Y_M_, Y_0_, and k were predicted by fitting Eqs. (3) to concentration-time data of intact protein as well as (4) and (5) to concentration-time data of degraded ^125^I-insulin and ^125^I-Aβ_40_, respectively. Afterwards, the whole model was fitted to heart radioactivity-time data, and the transfer rate constants *k*_*1*_, *k*_*2*_, and *k*_*0*_, were estimated using SAAM II (version 2.3, SAAM Institute, University of Washington, Seattle, WA). Residual plots were visually inspected to assess goodness of fit.

### Model validation and application

The developed model was validated using independent naïve datasets obtained from three young and three aged animals. To deconvolve plasma PK from the heart imaging data, the model was fitted to heart radioactivity-time data. After fixing the parameters *k*_*1*_, *k*_*2*_, and *k*_*0*_, plasma PK parameters were inferred by Bayesian estimation method, where prior estimates of population means and standard deviations of each parameter (A, B, α, β), obtained from our previous plasma PK studies (n=22 for ^125^I-insulin, n=27 for ^125^I-Aβ_40_), were employed. Heart tissue accumulation at the last time point (42 min) for each animal was also predicted and compared with observed heart radioactivity of both ^125^I-insulin and ^125^I-Aβ_40_ after transcardial perfusion.

The deconvolved plasma PK was used as input function to construct the Patlak plot by plotting,

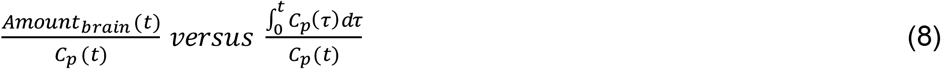

where *Amount*_*brain*_ (*t*) is brain radioactivity at time 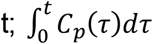 is plasma AUC from the time of injection; *C*_*p*_ (*t*) is the predicted plasma concentration of intact tracer at time t. The slope of the linear portion of the Patlak plot is referred to as the blood-to-brain influx clearance, Ki.

## Results

### Comparison of apparent heart PK and plasma PK of ^125^I-insulin and ^125^I-Aβ_40_

Following intravenous injection, the radioactivity signal in the heart detected by dynamic SPECT/CT imaging declined rapidly at early time points but became stable at the later phase **(Figure 1A, B)**. Radioactivity was quantified by ROI analysis and apparent heart concentration was obtained by taking the ratio of radioactivity and heart cavity volume. Notably, the apparent heart concentration was found to be higher than the observed plasma concentration of ^125^I-insulin **(Figure 1C)** or ^125^I-Aβ_40_ **(Figure 1D)** at all time points.

**Figure 1.**
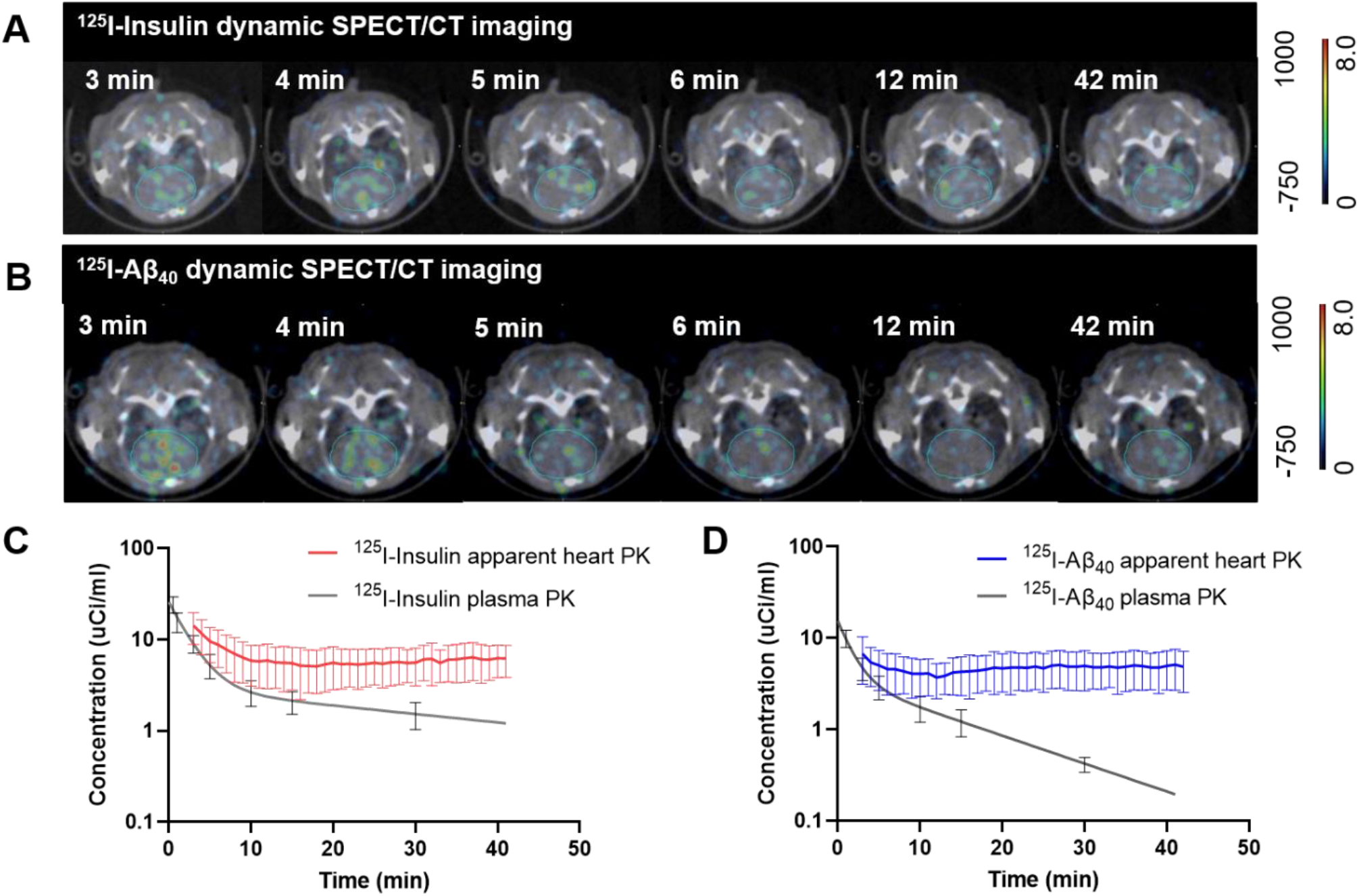
Apparent heart pharmacokinetics obtained by dynamic SPECT/CT imaging overestimates plasma pharmacokinetics of radiolabeled tracers ^125^I-insulin and ^125^I-Aβ_40_. Female B6SJLF1/J mice were intravenously injected with around 500 μCi ^125^I-insulin (n=8) or ^125^I-Aβ_40_ (n=9), and the heart of each animal was imaged up to 42 minutes by dynamic SPECT/CT. **(A-B)** Representative images of mice after injection with **(A)** ^125^I-insulin and **(B)** ^125^I-Aβ_40_ at 3 min, 4 min, 5 min, 6 min, 12 min, and 42 min and example regions of interest for data sampling are shown. **(C-D)** Comparison of apparent heart PK and plasma PK of **(C)** ^125^I-insulin and **(D)** ^125^I-Aβ_40_. Apparent heart PK was calculated as the ratio of heart radioactivity and heart cavity volume V_h_.

### Model fit of ^125^I-insulin and ^125^I-Aβ_40_ heart radioactivity-time profile

A three-compartment model was constructed, describing the radioactivity of intact and degraded radiolabeled proteins in plasma and their accumulation in the heart tissue **(Figure 2)**. Both intact ^125^I-insulin (**Figure 3A**) and ^125^I-Aβ_40_ (**Figure 3C**) exhibited biexponential disposition in plasma. The simple linear Eq. (4) adequately described the concentration-time data of degraded ^125^I-insulin (**Figure 3B**) whereas the nonlinear Eq. (5) was used to fit the concentration-time data of degraded ^125^I-Aβ_40_ (**Figure 3D**). Parameter estimates for all four forcing functions are summarized in **Table 1**. After fixing the parameters describing plasma disposition of the intact and degraded proteins, the model was fitted to the heart radioactivity-time data as determined by dynamic SPECT/CT imaging following intravenous injection of ^125^I-insulin (**Figure 4A**) or ^125^I-Aβ_40_ (**Figure 4B**). The model satisfactorily predicted heart tissue radioactivity, determined after transcardial perfusion, of both ^125^I-insulin and ^125^I-Aβ_40_ with relatively low prediction error **(Table 2)**. The goodness-of-fit plots for heart radioactivity data are displayed for ^125^I-insulin (**Supplementary figure 1)** and ^125^I-Aβ_40_ (**Supplementary figure 2)**. Individual predictions were in good agreement with observed values. Weighted residuals were found to be randomly distributed along the zero-ordinate line. Parameter estimates and their coefficients of variation (CV%) are presented in **Figure 4C, D**. Relatively low CV% values associated with the parameter estimates indicated goodness of the model fit.

**Table 1.** Parameter estimates of forcing functions describing plasma concentration-time profiles of intact proteins (A, B, α, β), degraded ^125^I-insulin (C_0_, C_1_) and ^125^I-Aβ_40_ (Y_M_, Y_0_, k) after IV bolus injection.

| $^{125}\text{I}$-insulin | | $^{125}\text{I}$-A$\beta_{40}$ | |
| --- | --- | --- | --- |
| Parameters | Estimates (Std. Error) | Parameters | Estimates (Std. Error) |
| **A** | 22.86 (2.46) | **A** | 11.95 (0.41) |
| **B** | 2.90 (0.57) | **B** | 3.50 (0.14) |
| **$\alpha$** | 0.43 (0.060) | **$\alpha$** | 0.62 (0.031) |
| **$\beta$** | 0.021 (0.0088) | **$\beta$** | 0.070 (0.002) |
| **$C_0$** | -0.27 (0.083) | **$Y_M$** | 3.714 (0.196) |
| **$C_1$** | 0.082 (0.0047) | **$Y_0$** | 0.463 (0.114) |
|  |  | **k** | 0.155 (0.027) |

**Table 2.**
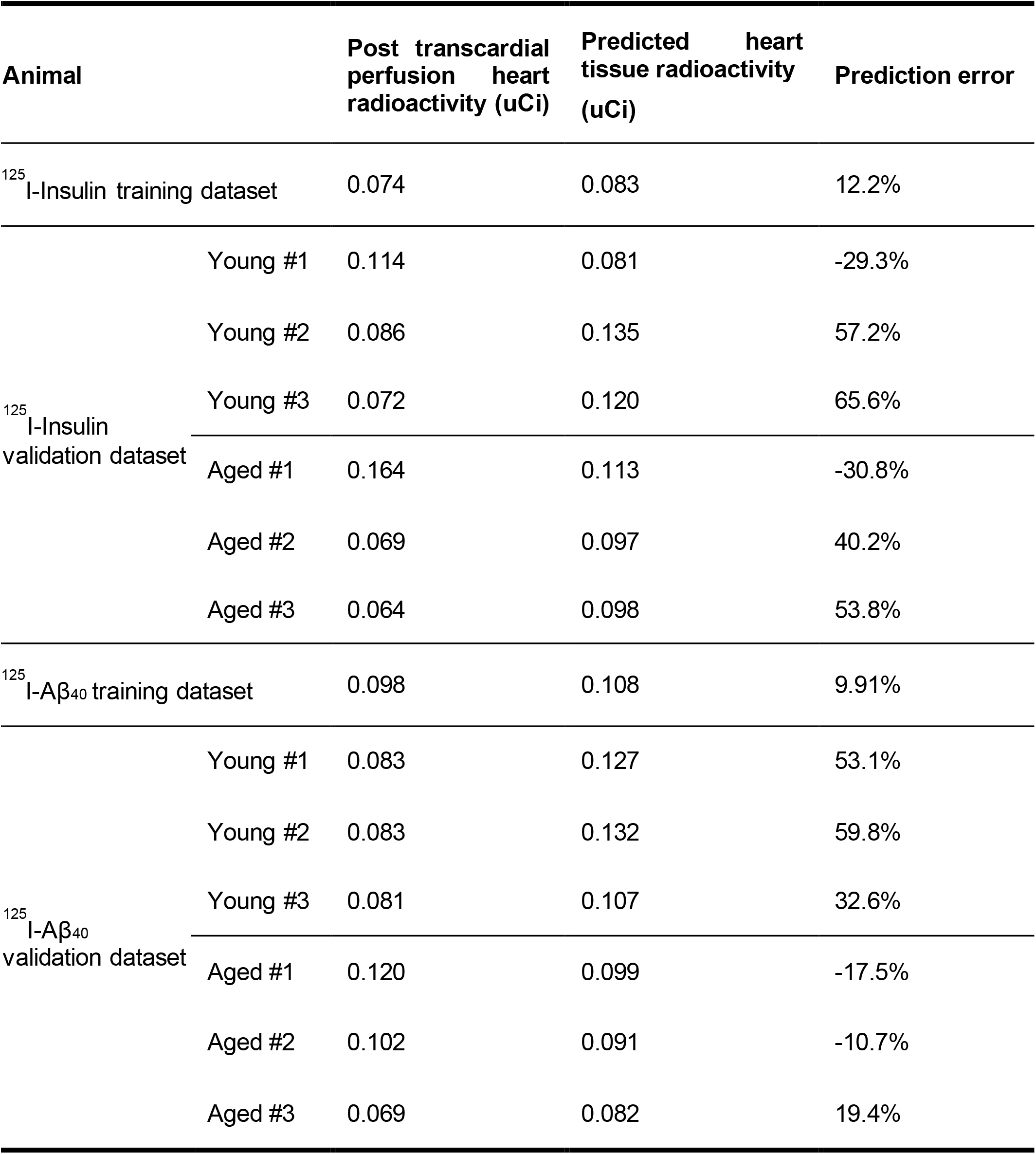
Comparison of model predicted heart accumulation and observed post transcardial perfusion heart radioactivity in training and validation datasets.

**Figure 2.**
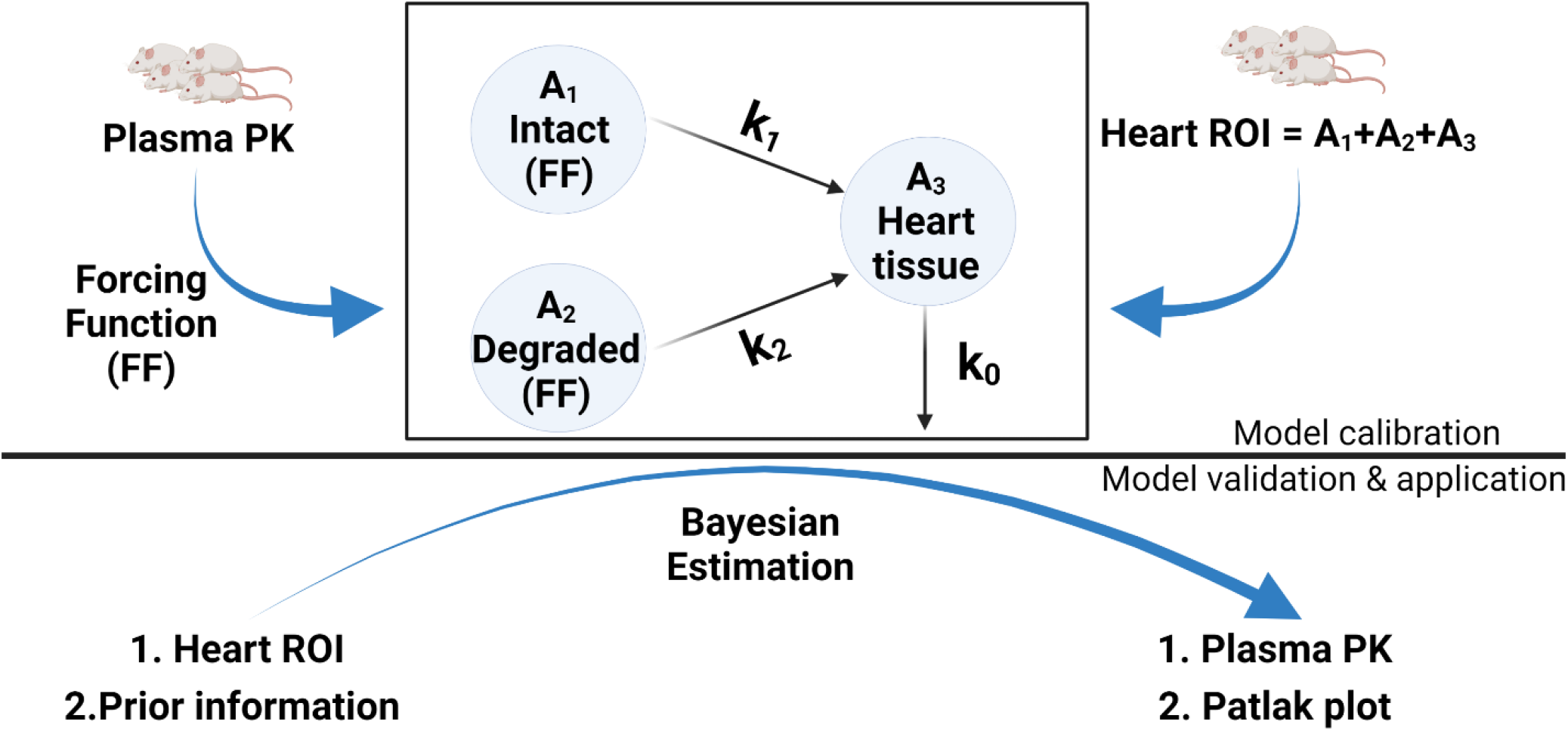
Model scheme and workflow. A three-compartment model was constructed including intact and degraded radiolabeled proteins in plasma (defined by forcing functions) and their accumulation in heart tissue. The model was fitted to the heart radioactivity-time data and transfer rate constants were estimated using SAAM II software. The model predictions were further verified using data from naïve animal group. Compartment 1 is intact protein (precipitate in TCA assay) and compartment 2 is degraded protein (supernatant in TCA assay) in the plasma. Compartment 3 is accumulation of radiotracers in the heart tissue. The parameters k_1_, k_2_, and k_0_ are first-order constants that describe transfer rates from compartments.

**Figure 3.**
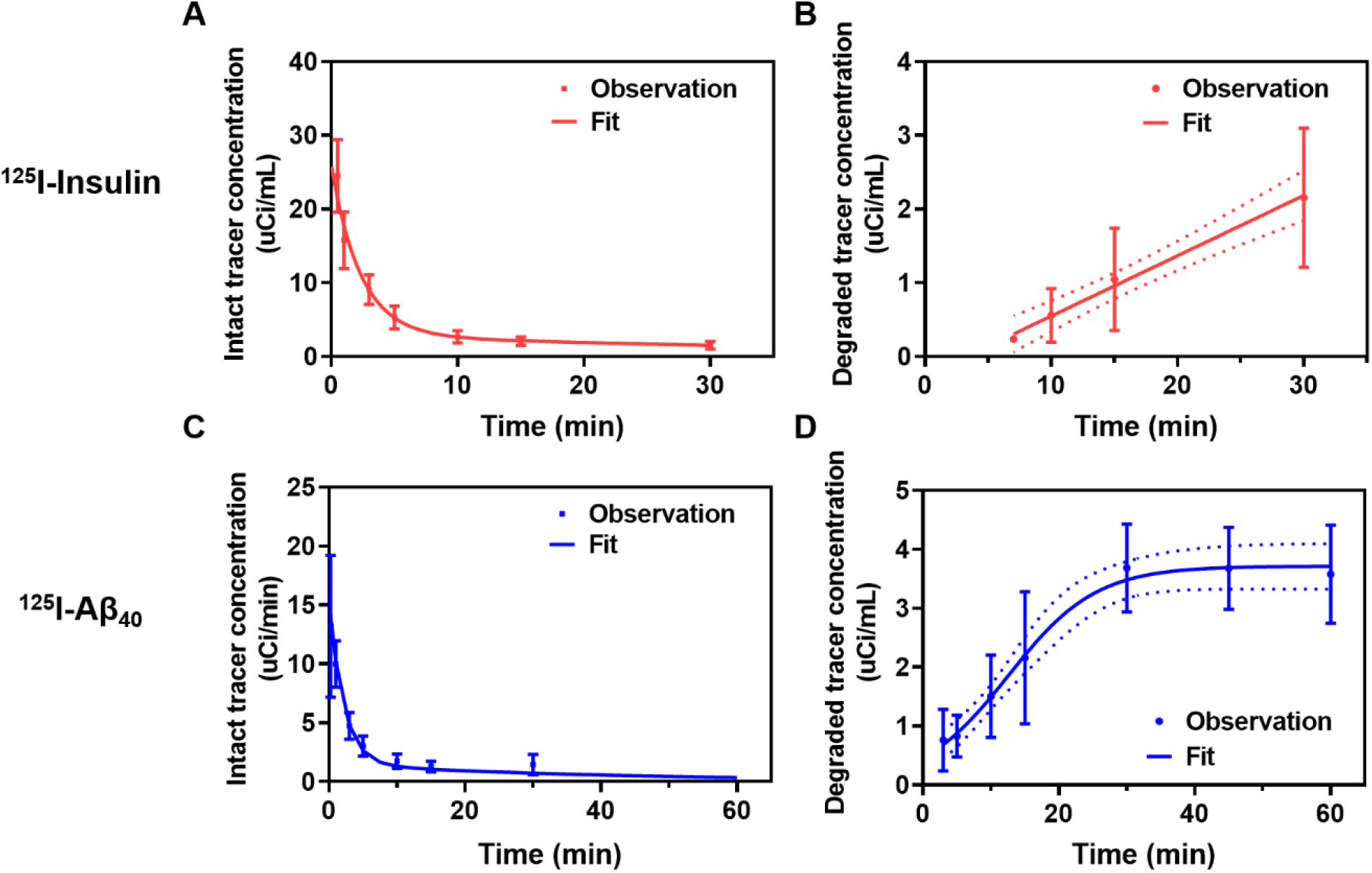
Forcing functions describing plasma concentration-time profile of intact and degraded ^125^I-insulin or ^125^I-Aβ_40_. Female B6SJLF1/J mice (n=12 for ^125^I-insulin, n=12 for ^125^I-Aβ_40_) were intravenously injected with 100uCi of ^125^I-insulin or ^125^I-Aβ_40._ Time series plasma samples were taken at regular intervals and are subjected to TCA precipitation. The plasma concentration in precipitate (intact) and supernatant (degraded) were determined by gamma count. Forcing functions were fitted to the plasma concentration-time data. **(A)** Intact ^125^I-insulin; **(B)** Degraded ^125^I-insulin; **(C)** Intact ^125^I-Aβ_40_; **(D)** Degraded ^125^I-Aβ_40_.

**Figure 4.**
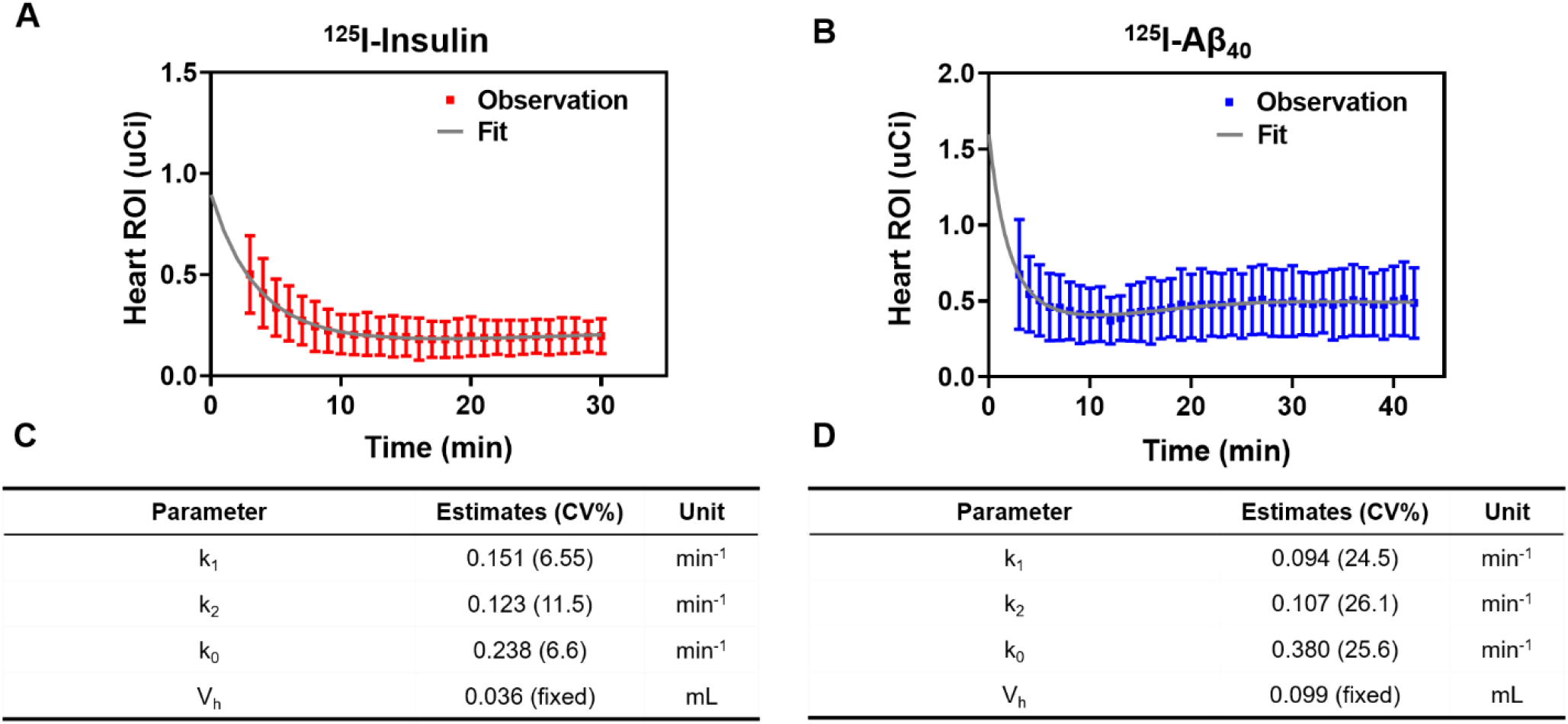
Model fit of the dynamic heart radioactivity data. The model shown in **Figure 2** was fitted to heart radioactivity-time data of **(A)** ^125^I-insulin and **(B)** ^125^I-Aβ_40_. Parameter estimates and coefficient of variation were presented in **(C)** ^125^I-insulin and **(D)** ^125^I-Aβ_40._ Volume of heart cavity *V*_*h*_ was determined separately as described in the materials and methods section and fixed during model fitting. *k*_*1*_, *k*_*2*_, and *k*_*0*_ are the first-order rate constants between the respective compartments.

### Model validation

The model thus developed was applied to deconvolve the plasma PK of both tracers from naïve dynamic heart imaging datasets. The model was found to adequately describe heart radioactivity-time datasets of both ^125^I-insulin (**Figure 5A**) and ^125^I-Aβ_40_ (**Figure 6A**) for six independent animals including three young (3-month-old) and three aged (24-month-old) mice. Further, plasma PK parameters including maximal concentration (C_max_), clearance (CL), and area under the curve (AUC) were predicted using the deconvolved plasma concentration-time datasets. In line with our previous report (L., Sharda et al. 2021 Dec 15), aged mice have significantly lower clearances compared to young mice for both ^125^I-insulin (**Figure 5C**) and ^125^I-Aβ_40_ (**Figure 6C**). The model was also validated by comparing model predicted tracer accumulation in heart tissue and observed heart radioactivity determined after transcardial perfusion. As shown in the **Table 2**, the predicted values were generally consistent with the observed heart radioactivity.

**Figure 5.**
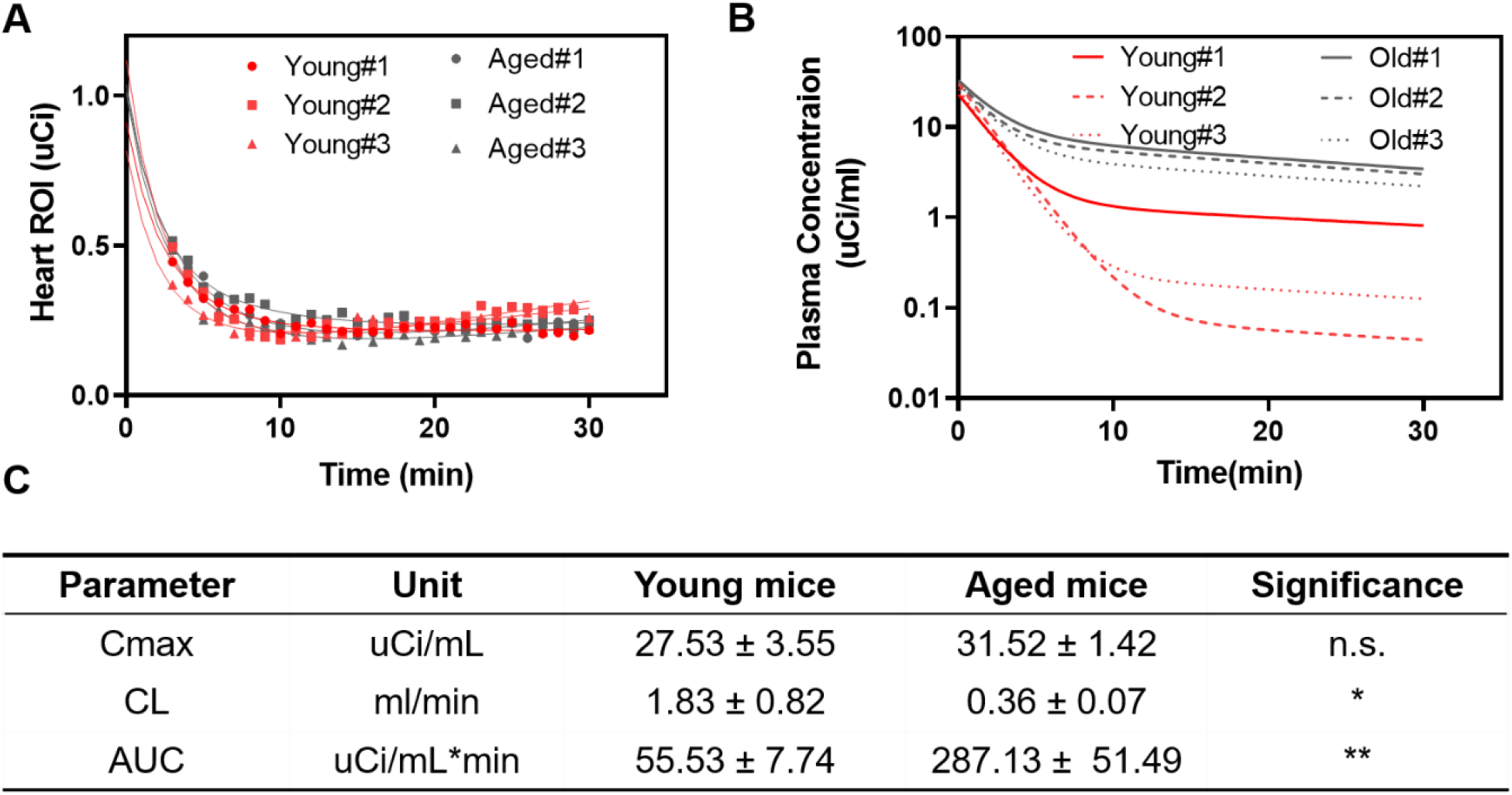
Prediction of ^125^I-insulin plasma PK in young and old mice from independent dynamic heart radioactivity datasets using the calibrated model. **(A)** Curve fitting of the dynamic heart radioactivity data. **(B)** Deconvolved plasma PK profile for each animal. **(C)** Comparison of various PK parameters derived from deconvolved plasma PK profile in young and old mice. *p<0.05, **p<0.01, student’s t-test.

**Figure 6.**
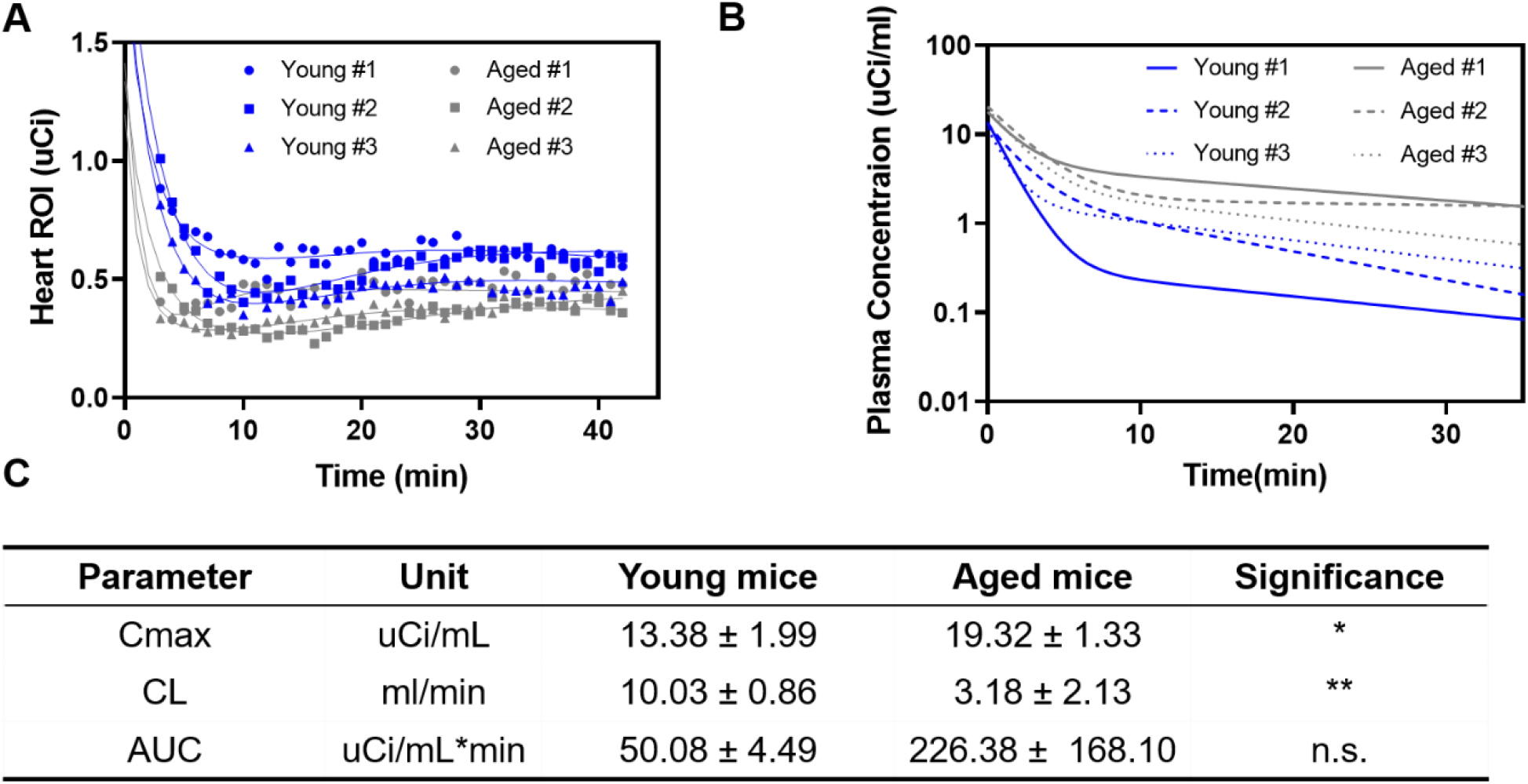
Prediction of ^125^I-Aβ_40_ plasma PK in young and old mice from independent dynamic heart radioactivity datasets using the calibrated model. **(A)** Curve fitting of the dynamic heart radioactivity data. **(B)** Deconvolved plasma PK profile for each animal. **(C)** Comparison of various PK parameters derived from deconvolved plasma PK profile in young and old mice. *p<0.05, **p<0.01, student’s t-test.

### Model application in Patlak plot analysis

Individually predicted plasma PK data from the developed model was employed to construct Patlak plots, and the predicted influx clearance (Ki) values were compared between young and aged mice. The representative Patlak plots for both ^125^I-insulin and ^125^I-Aβ_40_ were generated by three methods-using observed plasma PK (**Figure 7A, 8A**), deconvolved plasma PK (**Figure 7B, 8B**), or apparent heart PK (**Figure 7C, 8C**) as the input function. As summarized in **Figure 7D and Figure 8D**, methods employing observed plasma PK and deconvolved plasma PK were able to reproduce the Ki difference between young and aged mice as reported previously (L., Sharda et al. 2021 Dec 15). The Ki values predicted by these two methods were not significantly different. Using the deconvolved plasma PK, the Ki value of ^125^I-insulin was estimated to be 0.0018 ± 0.0008 ml/min in young mice and 0.0003 ± 0.0001 ml/min in aged mice, whereas the Ki values were predicted as 0.0013 ± 0.0005 ml/min in young mice and 0.0005 ± 0.0001 ml/min in aged mice using observed plasma PK. In the case of ^125^I-Aβ_40_, the estimated Ki value using deconvolved plasma PK was 0.0047 ± 0.0013 ml/min in young mice and 0.0025 ± 0.0004 ml/min in aged mice, whereas using observed plasma PK, the Ki value was estimated to be 0.0044 ± 0.0014 ml/min in young mice and 0.0018 ± 0.0003 ml/min in aged mice. However, analysis using apparent heart PK cannot capture age-dependent Ki differences. Moreover, the Ki values in aged mice were significantly lower than those analyzed using observed plasma PK.

**Figure 7.**
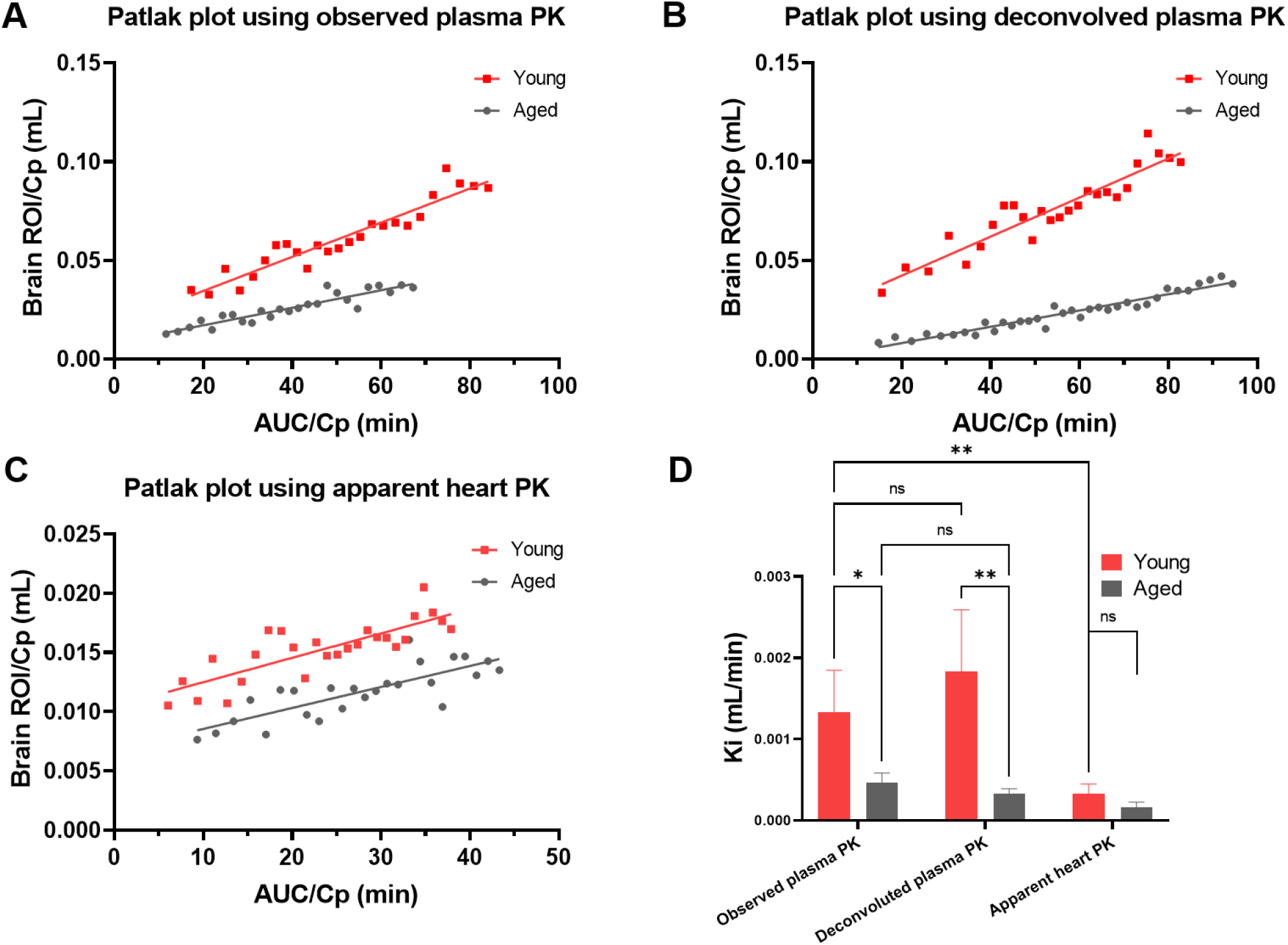
Patlak plot analysis of ^125^I-insulin in young and aged mice using different input functions. **(A-C)** Representative patlak plot using **(A)** observed plasma PK **(B)** deconvolved plasma PK and **(C)** apparent heart PK as input function. **(D)** Comparison of predicted blood-to-brain influx clearance values (Ki) from linear regression of respective Patlak plot. *p<0.05, **p<0.01, two-way ANOVA followed by multiple comparisons using two-stage set-up method of Benjamini, Krieger and Yekutieli.

**Figure 8.**
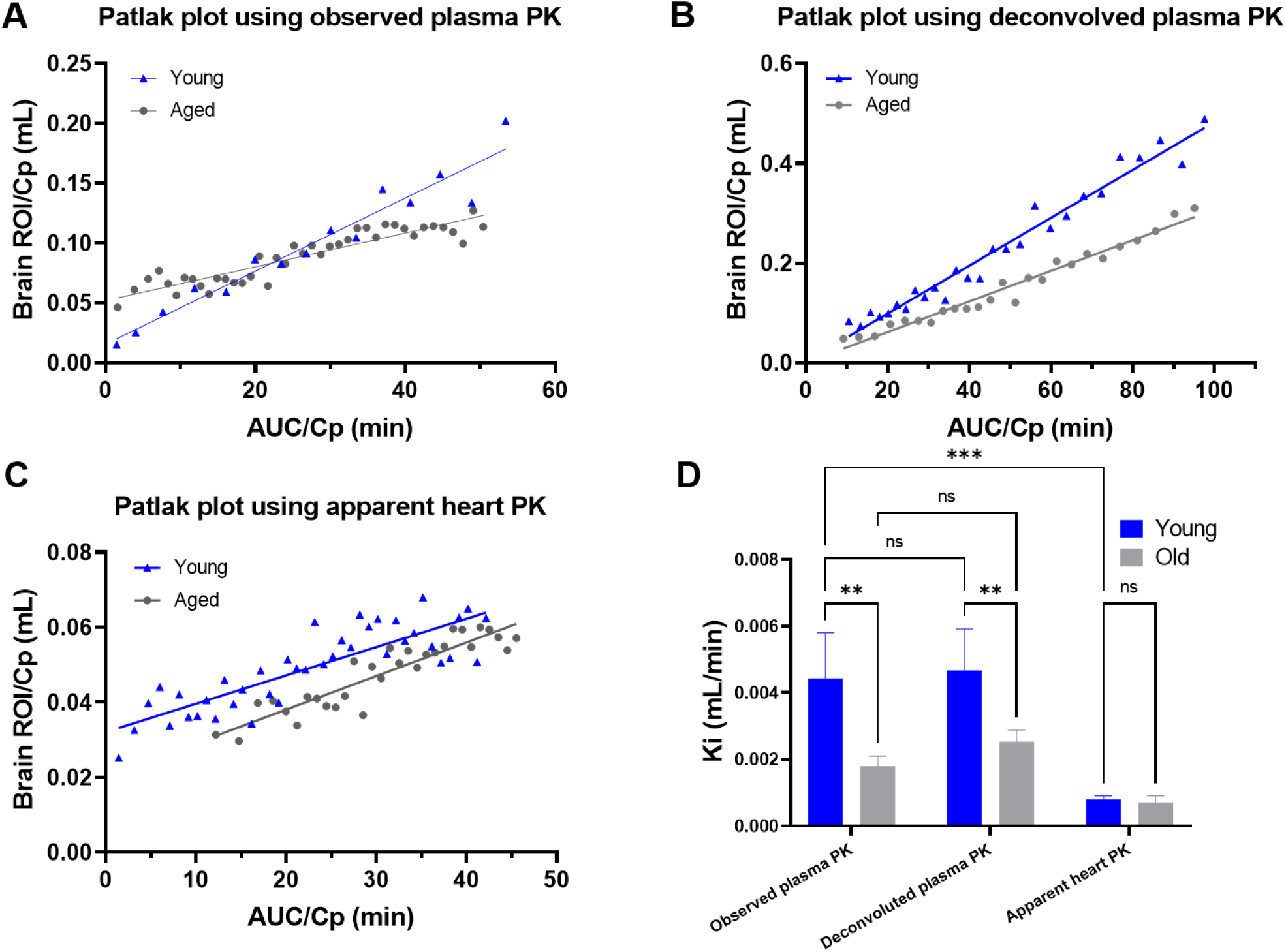
Patlak plot analysis of ^125^I-Aβ_40_ in young and aged mice using different input functions. **(A-C)** Representative patlak plot using **(A)** observed plasma PK **(B)** deconvolved plasma PK and **(C)** apparent heart PK as input function. **(D)** Comparison of predicted blood-to-brain influx clearance values (Ki) from linear regression of respective Patlak plot. **p<0.01, ***p<0.001, two-way ANOVA followed by multiple comparisons using two-stage set-up method of Benjamini, Krieger and Yekutieli.

## Discussion

In vivo small animal imaging is widely used to assess bio-distribution of drugs, where an imaging probe is administered to visualize the tissue of interest (Rudin and Weissleder 2003 Feb, Gomes, Abrunhosa et al. 2011, Ding and Wu 2012). Various imaging modalities have been developed including optical imaging (OI), ultrasonography (US), computed tomography (CT), magnetic resonance imaging (MRI), and nuclear imaging, involving positron emission tomography (PET) and single photon emission computed tomography (SPECT). Dynamic SPECT/CT imaging is commonly employed to characterize the uptake kinetics of radiolabeled tracers into tissue of interest by graphical analysis such as the Patlak plot. In this case, tissue radioactivity as quantified from dynamic imaging and plasma PK data as input function are integrated through the Patlak equation and the influx rate is estimated by linear regression (Gjedde 1982, Patlak, Blasberg et al. 1983, C.S., Blasberg et al. 1983 Mar). Using this method, we have assessed the blood-to-brain influx rate of ^125^I-insulin, ^125^I-Aβ_40_ and ^125^I-Aβ_42_ in mouse models under different pathophysiological conditions (Swaminathan, Ahlschwede et al. 2018, Sharda, Ahlschwede et al. 2021). However, in those studies, the plasma PK and dynamic SPECT/CT imaging data were acquired from separate animal cohorts due to the infeasibility of simultaneous dynamic imaging and plasma sampling. Consequently, extra effort was made to conduct plasma PK studies and only the group-averaged plasma concentration-time data instead of that obtained from individual mice was used as the input function for Patlak analysis. Although longitudinal imaging data of heart has been used as a surrogate for plasma PK of various tracers (Green, Nguyen et al. 2004, Giddabasappa, Gupta et al. 2016), our results demonstrate that apparent heart PK overestimates ^125^I-Insulin **(Figure 1C)** and ^125^I-Aβ_40_ **(Figure 1D)** plasma levels, with larger error in the terminal phase. This observed overprediction was most likely due to substantial accumulation of these tracers, especially Aβ, in the heart tissue. Heart accumulation of amyloid forming proteins has been widely reported in the literature to cause cardiac amyloidosis (Martinez-Naharro, Hawkins et al. 2018).

In the present study, we developed a three-compartment pharmacokinetic model to deconvolve the plasma PK of radiotracers, ^125^I-insulin and ^125^I-Aβ_40_, from their heart dynamic SPECT/CT imaging data. This procedure provides a novel approach to characterizing plasma PK of drugs by non-invasive imaging. Further, it offers benefits in reduction of number of animals needed and more accurate estimation of Patlak parameters for describing uptake rate of macromolecules into tissue of interest under various pathophysiological conditions.

The PK model enables separation of the vascular and extravascular contributions to the total radioactivity in the heart. Additionally, vascular radioactivity could consist of the radioactivity of intact protein (compartment 1) and the degradation products that carry the radiolabel (compartment 2) (**Figure 2**). Both insulin (Duckworth, Bennett et al. 1998) and Aβ (Xiang, Bu et al. 2015) metabolites have been shown to circulate in plasma and each may have distinct disposition profiles from their parent molecules. Therefore, incorporating a separate compartment representing degraded metabolites could significantly improve the predictive ability of a model describing their disposition. The model was calibrated stepwise against plasma concentration-time data of intact and degraded protein tracers, and the heart radioactivity-time profile as determined by SPECT/CT dynamic imaging.

Vascular radioactivity of intact and degraded protein was described by a forcing function which is the product of plasma concentration and volume of the heart cavity. Only plasma instead of whole blood concentration was considered because neither insulin nor Aβ exhibit significant uptake by blood cells based on our previous studies. Therefore, vascular radioactivity was assumed to be substantially contributed by tracers in plasma. The forcing function is a useful tool in pharmacokinetic modeling and is generally used as a model input to simulate PK profiles of drugs following different dosing regimens or under various pathophysiological conditions (Zhang, Wang et al. 2012, Hof and Bridge 2021). In this study, a biexponential equation, which adequately describes the concentration-time data of both intact ^125^I-insulin and ^125^I-Aβ_40_ peptides, was used as the forcing function (**Figure 3A, C**). These results were consistent with previous reports where two-compartment models were used to fit plasma PK data of these two peptides in mouse (Kandimalla, Curran et al. 2005, L., Sharda et al. 2021 Dec 15).

Unlike intact proteins, the concentrations of degradation products, characterized by supernatant counts after TCA precipitation, were found to increase over time for both ^125^I-insulin and ^125^I-Aβ_40_, possibly due to inefficient clearance of metabolites from systemic circulation. Notably, within the time frame of the experiment, increase of degraded ^125^I-insulin was found to be linear (**Figure 3B)** whereas ^125^I-Aβ_40_ metabolite appearance was nonlinear and reached a plateau at later time points **(Figure 3D)**. Interestingly, the intercept (C_0_) of the linear equation describing ^125^I-Insulin metabolite concentration-time profile was found to be negative **(Table 1)**. One possible reason for this outcomeis that there is a lag time for ^125^I-insulin metabolite to appear in the systemic circulation.

Volume of heart cavity (V_h_) was separately determined using post-perfusion radioactivity data obtained from heart dynamic imaging. The V_h_ value for ^125^I-insulin (35 μL) was found to be lower than that of ^125^I-Aβ_40_ (99 μL). One plausible explanation is the facile association of ^125^I-Aβ_40_ to the cavity wall and/or to the heart tissue that could be washed off during perfusion. Thus, greater loss of post-perfusion radioactivity may apparently increase the estimate of heart cavity volume. This hypothesis was supported by literature reports pertaining to Aβ disposition in the heart. For instance, Aβ receptors such as receptor for advanced glycation end products (RAGE) and lipoprotein receptor-related protein 1 (LRP1) are shown to be expressed in various heart cells, including endothelial cells, cardiomyocytes, and fibroblasts (Ramasamy and Schmidt 2012) (Potere, Del Buono et al. 2019). Further, Aβ aggregates have been found in hearts of AD patients and affect myocardial function (Troncone, Luciani et al. 2016, Martinez-Naharro, Hawkins et al. 2018), which is implicated in the so-called “cardiogenic dementia” that links dementia to cardiovascular disease (T.R., Milne et al. 1981 Oct 3). Although insulin receptors are ubiquitously expressed and are also highly enriched in the heart (Riehle and Abel 2016, Abel 2021 Jul 1), deposition of insulin in heart tissue has not been reported. Nevertheless, the calculated V_h_ values were comparable to reported anatomical volume of the heart cavity in mice. Previous studies have reported that the stroke volume, which pertains to the volume of blood pumped out of the left ventricle, in mice ranges from 0.025 to 0.04 ml (Barbee, Perry et al. 1992). Considering an ejection fraction of around 50% in anesthetized mice (Vinhas, Araujo et al. 2013), the total blood volume of heart should be around 50 to 80 μL. After successfully identifying forcing functions of plasma PK and cavity volume, these parameters were fixed and the transfer rate constants (k_1_, k_2_, k_0_) were inferred by fitting the model to the heart radioactivity-time data. As shown in **Figure 4A, C**, the fitted heart radioactivity of both ^125^I-insulin and ^125^I-Aβ_40_ were in excellent agreement with observed data, which was also indicated by the goodness-of-fit plots (**Supplementary figure 1 and 2)**. Moreover, all the rate constant estimates were shown to have low CV%, confirming the precision of the parameter estimation from our data.

Upon identifying all the model parameters, the model was applied to deconvolve plasma PK profiles of ^125^I-insulin and ^125^I-Aβ_40_ from independent heart radioactivity-time datasets. To further validate the predictive ability of our model, we used heart imaging datasets obtained from both young and aged mice and the model is expected to reproduce age-dependent plasma PK changes as we reported previously. Specifically, increased plasma exposure (AUC) with aging was observed for both ^125^I-insulin and ^125^I-Aβ_40_ (L., Sharda et al. 2021 Dec 15). For deconvolution, the developed model was fitted to heart radioactivity-time data after fixing V_h_ as well as transfer rate constants (k_1_, k_2_, k_0_) and leaving the plasma PK parameters adjustable. To avoid over-parameterization, the Bayesian estimation method, widely used in population PK/PD analysis (Callegari, Caumo et al. 2002, Bland, Pai et al. 2018), was used during curve fitting. As shown in **Figures 5A and 6A**, the fitted dynamic heart imaging data agreed well with the observed data obtained from all six mice for both tracers. Moreover, the predicted plasma PK profile demonstrated significantly decreased clearance and thus increased AUC in aged mice compared to young mice for both tracers, which is consistent with our previous observation (L., Sharda et al. 2021 Dec 15). Nevertheless, these results indicated that our modeling approach could successfully deconvolve plasma PK of ^125^I-insulin and ^125^I-Aβ_40_ from heart imaging data under different pathophysiological conditions and capture age-dependent plasma PK changes.

The model predictive ability was further verified by comparing the predicted heart tissue accumulation at the last time point (42 minutes) with observed post-transcardial perfusion heart radioactivity obtained from SPECT/CT imaging. The prediction error was found to be around or less than 50% for most of independent validation animals. However, for 8 out of 12 validation animals, our model appeared to overestimate heart tissue radioactivity **(Table 2)**, most likely due to loss of tracers from heart tissue during perfusion as described above, which could yield lower radioactivity than the model prediction.

The modeling approach for deconvolution of plasma PK from dynamic heart imaging data is very useful because: a) plasma PK analysis and imaging studies can be conducted in a single cohort of animals, substantially reducing the number of animals and conserving resources; b) it improves the accuracy and precision of Patlak plot parameter estimates since the plasma PK and dynamic imaging data are obtained from the same mouse. Although bulk heart imaging data has been found to agree with plasma PK of probes that are restricted to the vascular space (Bao, Vasquez et al. 2019), it is not applicable to molecules that are substantially taken up by the heart tissue. For example, a previous study reported that heart ROI time-activity curves (TACs) can overestimate the actual blood concentration of ^18^F-FDG especially at later time points after injection. Consequently, the brain influx parameter estimated using heart ROI TAC shows a larger error (Green, S.S. et al. 1998 Apr). Our modeling approach could address this issue by separating the vascular and extravascular radioactivity of the whole heart and serves as a better surrogate for plasma PK. Hence, we used plasma PK profiles resulting from this modeling approach to generate Patlak plots in both young and old mice. The results indicated that blood-to-brain influx clearance, Ki, of ^125^I-insulin and ^125^I-Aβ_40_ decreased significantly with aging, which is consistent with our previous publications (L., Sharda et al. 2021 Dec 15). In contrast, when apparent heart PK was used as the input function, the difference between young and old mice was lost due to inaccurate estimation of real plasma PK. These results established the feasibility of our modeling method to investigate blood-to-tissue influx rate without conducting extra plasma PK studies. Notably, the brain and heart imaging were obtained from the same animal so that the estimated Ki is expected to be more accurate. Successful application of the modeling method established in this study, will allow us to infer distribution kinetics of other tracers in both health and disease. Indeed, we have employed this approach to evaluate blood-to-brain influx of a newly developed PET probe ^68^Ga-NOTA-insulin in wild type and Alzheimer’s transgenic mice (Taubel, Nelson et al.).

In conclusion, we have developed a simple compartmental modeling approach to deconvolve plasma PK of radiotracers from non-invasive dynamic heart imaging data obtained from mice. Successful prediction of age-dependent changes in the plasma pharmacokinetics of ^125^I-insulin and ^125^I-Aβ_40_ confirms its utility in assessing systemic disposition of various macromolecular tracers. Moreover, application of this method to generate Patlak plots allows for accurate prediction of tissue influx kinetics of radiotracers, especially when simultaneous plasma sampling is not feasible.

## Acknowledgement

The authors would like to acknowledge Dr. Ronald J. Sawchuk for his hands-on instructions on the SAAM II software. The graphic abstract/Figure 2 was generated using BioRender.

## Funding Statement

This work was supported by the Minnesota Partnership for Biotechnology and Medical Genomics (MNP#15.31) and National Institutes of Health/National Institute of Neurological Disorders and Stroke (R01NS125437).

## Conflict of Interest Statement

Dr. Lowe reported consulting for Bayer Schering Pharma, Piramal Life Sciences, Life Molecular Imaging, Eisai Inc., AVID Radiopharmaceuticals, and Merck Research and receiving research support from GE Healthcare, Siemens Molecular Imaging, AVID Radiopharmaceuticals and the NIH (NIA, NCI). The other authors declared no potential conflicts of interests with respect to the research, authorship, and/or publication of this article.

## Author Contributions

*Participated in research design:* Z Wang, L Wang, Ebbini, Kandimalla

*Conducted experiments:* Z Wang, L Wang, Curran, Vernon

*Contributed analytic tools:* Min, Lowe

*Performed data analysis:* Z Wang, L Wang

*Wrote or contributed to the writing of the manuscript:* All authors

## Notes

### Competing Interest Statement

The authors have declared no competing interest.

