## Supplementary figures for "Deconvolution of plasma pharmacokinetics from dynamic heart imaging data obtained by SPECT/CT imaging"

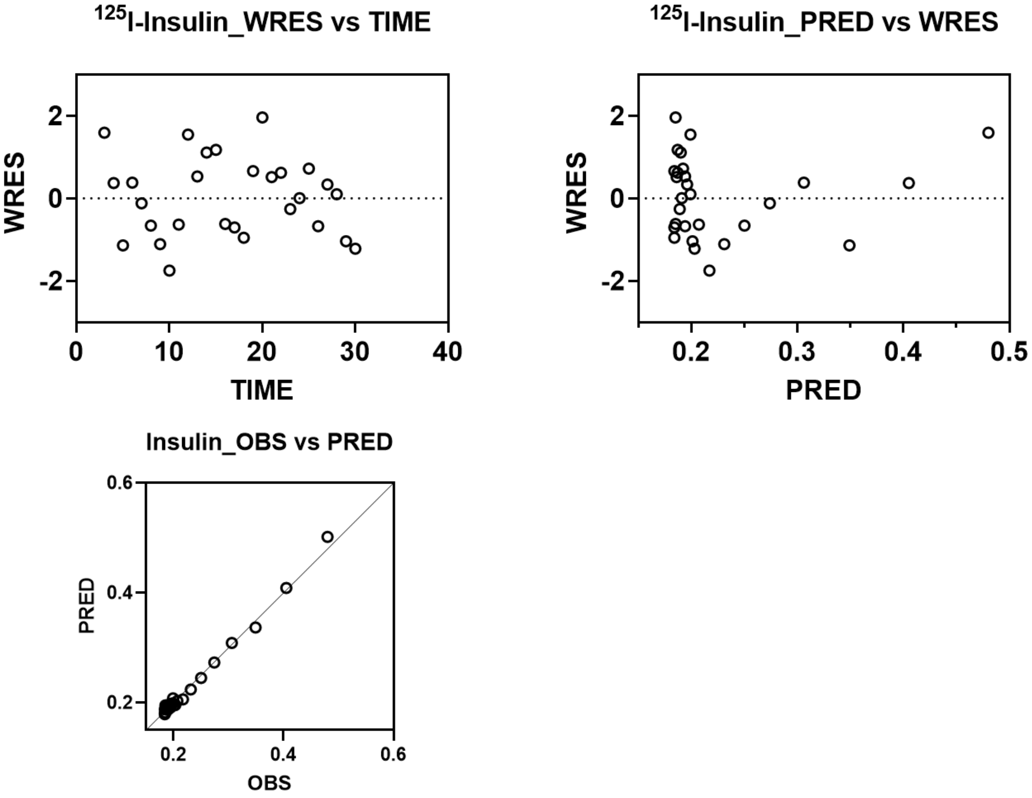


**Supplementary figure 1. Diagnostic plots for $^{125}$I-insulin model fitting.**

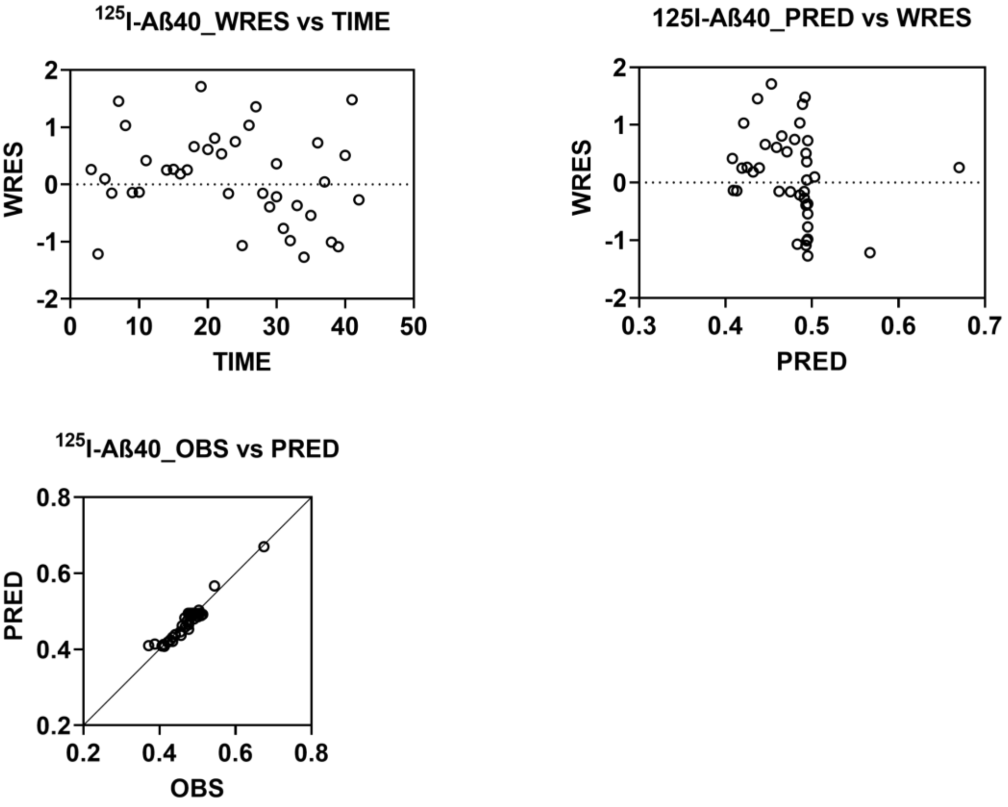


**Supplementary figure 2. Diagnostic plots for $^{125}$I-Aβ40 model fitting.**
